# Visual Analysis of Cell Death in Bladder Cancer — A Bibliometrics-Based Comprehensive Study

**DOI:** 10.1101/2025.08.10.669556

**Authors:** Zhi-wen Cao, Yue-yi Chen, Nan-xi Fu, Fu-rui Fu, Sen-jie Shi, Hong-yu Wang, Chuang-long Xu

## Abstract

Cell death mechanisms offer therapeutic strategies for bladder cancer, yet lack comprehensive bibliometric analysis. Map global research trends and hotspots in bladder cancer cell death mechanisms via bibliometrics. We analyzed 5,392 publications (1991–2024) from Web of Science using VOSviewer (co- authorship/keyword clustering), CiteSpace (citation bursts), and GraphPad Prism (statistics). Metrics included: (1) Temporal trends, (2) Country/institution contributions, (3) Journal impact, (4) Citation dynamics, (5) Collaboration networks, (6) Conceptual hotspots. China and the US led research output. Top institutions:University of Texas System (USA; 178 publications), UTMD Anderson Cancer Center (USA; 123), Nanjing Medical University (China; 122), Journal of Urology had the highest output (106 publications); Cancer Research (IF:12.5) the highest impact. Kim Wun-Jae was the most productive author (37 articles); Jemal A the most co-cited (446 citations). Keyword and citation analyses revealed emerging integration of cell death mechanisms with immunotherapy (IT) and photodynamic therapy (PDT) to overcome chemoresistance. This study delineates the evolution of bladder cancer cell death research and identifies IT/PDT as promising resistance-overcoming strategies grounded in targeted cell death pathways.

## 1 Introduction

Bladder cancer (BC) constitutes the most frequent urogenital malignancy. GLOBOCAN 2022 reports BC as the ninth most common cancer worldwide and the thirteenth leading cause of cancer-related mortality, while epidemiological studies consistently demonstrate a 3–4-fold higher incidence in males compared to females **Error! Reference source not found.**. Urothelial carcinoma (UC) constitutes the predominant histological subtype of bladder tumors. Histopathological classification is based on detrusor muscle invasion status: non-muscle-invasive bladder cancer (NMIBC) refers to tumors confined to the mucosal epithelium or lamina propria, whereas muscle-invasive bladder cancer (MIBC) denotes invasion into the muscularis propria layer **Error! Reference source not found.**. Among these, NMIBC accounts for approximately 75% of BC cases, while MIBC comprises the remaining 25%. Studies report alarmingly high 5-year recurrence and progression rates for NMIBC— up to 78% and 45%, respectively—which complicate clinical management and elevate recurrence risks. For intermediate- to high-risk NMIBC, the cornerstone initial therapy involves transurethral resection of bladder tumor (TURBT) followed by adjuvant intravesical Bacillus Calmette-Guérin (BCG). Notably, BCG immunotherapy remains the only established, widely adopted conservative treatment proven to reduce progression risk in high-risk NMIBC **Error! Reference source not found.Error! Reference source not found.**. Conversely, MIBC typically presents with high-grade histology, locally advanced disease, or metastases, necessitating cisplatin-based neoadjuvant chemotherapy and radical cystectomy, often supplemented by biomarker-guided immune checkpoint inhibitors or targeted combination therapies **Error! Reference source not found.Error! Reference source not found.**. Although current strategies (e.g., TURBT, radical cystectomy, chemotherapy, and immunotherapy) form the therapeutic backbone, persistent challenges include high recurrence rates and treatment resistance in advanced/metastatic BC. Consequently, elucidating pathogenic mechanisms and identifying actionable therapeutic targets through fundamental research is critically needed to improve clinical outcomes.

As a genitourinary malignancy, BC pathogenesis is closely linked to cell death mechanisms. Diverse forms of cell death—including apoptosis (regulated by factors such as Bcl-2 and survivin), pyroptosis, and ferroptosis—play critical roles in BC initiation, progression, and treatment response. Dysregulation of apoptosis-related gene expression is a well-established key factor in the development and advancement of numerous cancers **Error! Reference source not found.**. Pyroptosis, identified as a distinct form of programmed cell death (PCD), demonstrates a close association with urothelial cell damage in BC development; consequently, several compounds can mitigate BC progression by inhibiting pyroptosis **Error! Reference source not found.**. Compared to normal cells, tumor cells require increased iron for malignant proliferation, implying heightened tumor susceptibility to ferroptosis. Crucially, ferroptosis contributes significantly to BC pathogenesis and can induce death in apoptosis-resistant tumor cells**Error! Reference source not found.**. Therefore, these cell death mechanisms are fundamentally implicated in BC. Although substantial clinical research on bladder cancer cell death has been documented and significant progress achieved, current studies remain fragmented. Critically, a systematic analysis delineating the fields developmental trajectory, core research foci, and emerging frontiers is lacking. To address this gap, we conducted a comprehensive bibliometric study.

Bibliometrics employs statistical methods to quantitatively analyze scholarly publications, systematically mapping research trajectories and intellectual landscapes. Through mathematical techniques, it elucidates academic contributions—including affiliations, journals, authors, citations, and keywords—while integrating data visualization to assess literature volume, quality, impact, and structural relationships **Error! Reference source not found.Error! Reference source not found.**. This approach provides an ideal framework for identifying influential research, emerging hotspots, and global trends, thereby generating actionable insights for future investigations. Applying this methodology, our study integrates statistical analysis with advanced visualization tools (e.g., co-citation networks, keyword clustering) to establish the first bibliometric overview of global research on cell death mechanisms in bladder cancer. We further identify evolving research trends and future directions. The findings provide researchers and clinicians with a definitive roadmap of the fields evolution and highlight priority areas for targeted exploration.

## 2 Data and Methods

### 2.1 Data source and retrieval strategy

The WoSCC, a premier multidisciplinary database indexing high-impact, peer-reviewed journals, was selected for this bibliometric analysis. Its rigorous curation ensures the inclusion of high-quality bladder cancer research, aligning with our objective to provide a reliable and comprehensive field assessment. On December 31, 2024, we systematically retrieved publications using the following search strategy: (Cell Death Terms): TS=("Cell Death" OR "Death, Cell" OR Apoptosis OR "Classical Apoptosis" OR "Classic Apoptosis" OR "Programmed Cell Death, Type I" OR "Extrinsic Pathway Apoptosis" OR "Intrinsic Pathway Apoptosis" OR "Programmed Cell Death" OR "Caspase-Dependent Apoptosis" OR Pyroptosis OR "Inflammatory Apoptosis" OR "Pyroptotic Cell Death" OR "Caspase-1 Dependent Cell Death" OR Necroptosis OR Ferroptosis OR Disulfidptosis) AND (Bladder Cancer Terms): TS=("bladder cancer*"* OR "Bladder Neoplasm*"* OR "Bladder Tumor*"* OR "Cancer of Bladder*"* OR "Urinary Bladder Cancer*"). Inclusion Criteria: English-language original research, reviews, experimental studies, and clinical trials. Exclusion Criteria: Conference abstracts, book chapters, and non-peer-reviewed publications. The initial search yielded 5,880 records. After duplicate removal using EndNote followed by manual verification, two independent authors screened titles/abstracts. Publications unrelated to bladder cancer cell death mechanisms were excluded. Discrepancies were resolved through discussion with a third author. This process resulted in 5,392 articles for final analysis (Fig 1).

**Fig 1.**
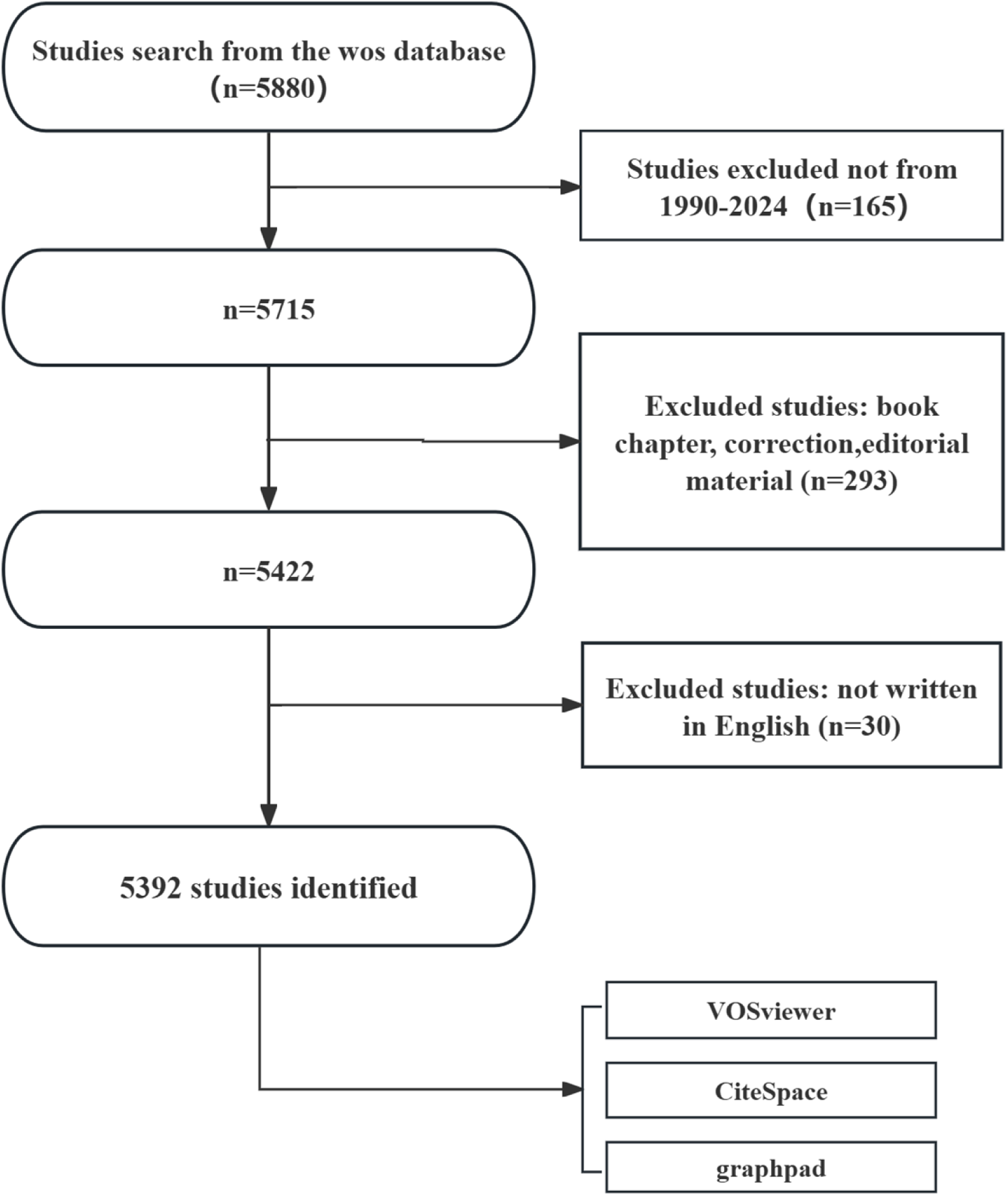
Flowchart of literature search

### 2.2 Data analysis and visualization

This study employed GraphPad Prism, CiteSpace, and VOSviewer for comprehensive data analysis and visualization, focusing on annual publication counts, national/regional publication trends, proportional distributions, and the generation of scientific knowledge maps. Specifically, GraphPad Prism was utilized to calculate and visualize publication volumes by year, publication trends across regions/countries, and proportional research distributions. VOSviewer facilitated the analysis of collaborations among countries/regions, institutions, and authors, as well as keyword clustering. CiteSpace was employed to analyze citation bursts, keyword co-occurrence patterns, and cited reference networks, thereby identifying emerging research frontiers and intellectual bases within the field. In the resulting visualizations, each node represents an element (e.g., country, institution, keyword); node size corresponds to publication count (for countries/institutions) or keyword frequency, while connecting lines denote collaboration strength or keyword co-occurrence frequency, with colors differentiating clusters or temporal periods. Furthermore, CiteSpace enabled the examination of citation burst evolution and keyword emergence trends over time, thereby providing insights into the developmental trajectory of the research field.

## 3 Results

### 3.1 Global trend in publication outputs andcitations

Analysis of the WoSCC revealed a total of 5,392 publications on bladder cancer cell death indexed as of December 31, 2024. These publications originated from 654 institutions across 95 countries/regions, were published in 1,235 journals, and involved contributions from 1,693 authors. Annual publication trends are presented in Fig 2. The field exhibited a consistent upward trajectory, with an average annual growth rate of 14.44% and a peak output of [specify number] publications in 2022. This trend signifies a substantial and progressive increase in global research activity focused on bladder cancer cell death. Furthermore, the pronounced acceleration in recent years strongly suggests that bladder cancer cell death will remain a prominent and actively investigated area with significant research potential in the foreseeable future.

**Fig 2.**
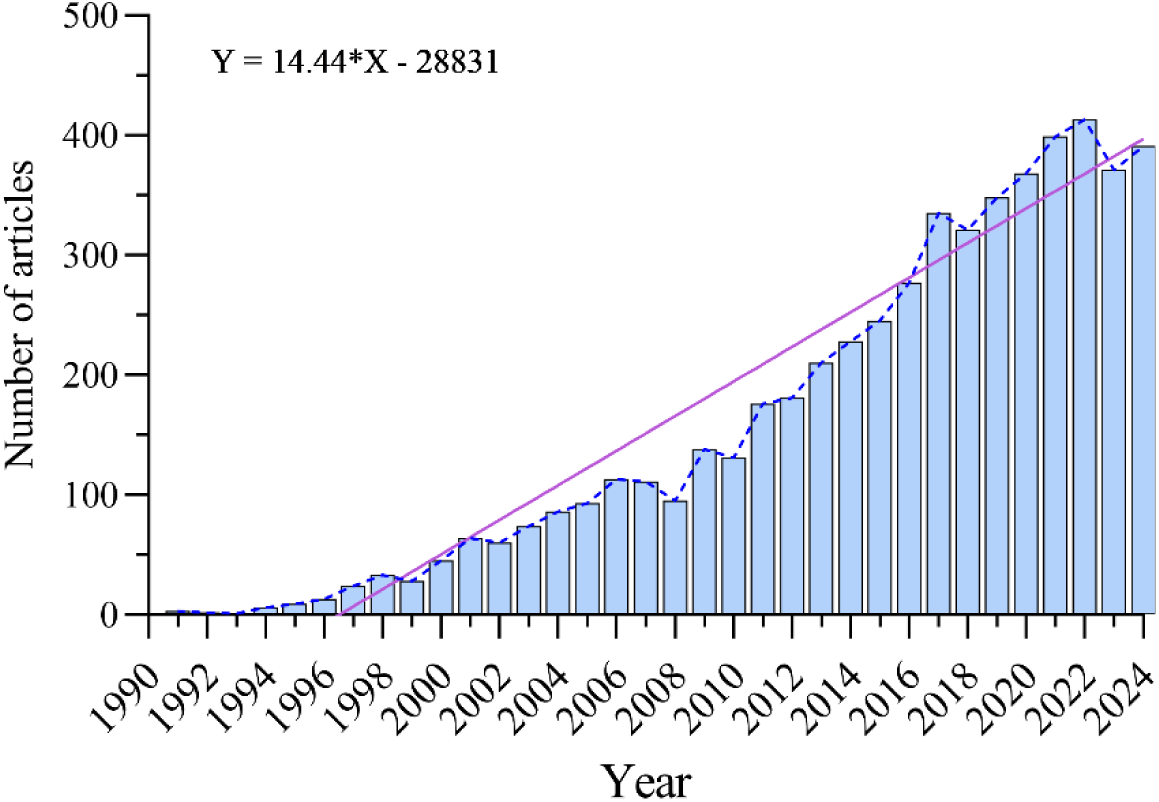
Annual volume of publications

### 3.2 Distribution of countries/regions

A total of 95 countries/regions have contributed to bladder cancer cell death research. Figs 3A and 3B illustrate the annual publication outputs of the top 10 productive countries/regions over the past decade. The top five contributors were China, the United States, Japan, Germany, and South Korea. China accounted for 45.38% of total publications, significantly exceeding the combined output of all other nations. Among the top 10 countries/regions, Chinese publications accumulated 62,119 citations (Table 1)—substantially higher than all others—with a citations-per-publication ratio of 25.39 (ranking 7th globally), indicating consistently high research quality. The United States ranked second in publications (1,093 papers), citations (54,042), and citations-per-publication ratio (49.45), reflecting strong citation influence. Notably, the United Kingdom, though ranking 7th in publications (152 papers), achieved the highest citations-per-publication ratio (53.63), demonstrating exceptional impact and scholarly quality. Both the collaborative network (Fig 3C) and geospatial coordination map (Fig 3D) underscore the dominant scholarly leadership of China and the United States, as their disproportionately large node sizes signify superior productivity and academic engagement in this field.

**Fig 3.**
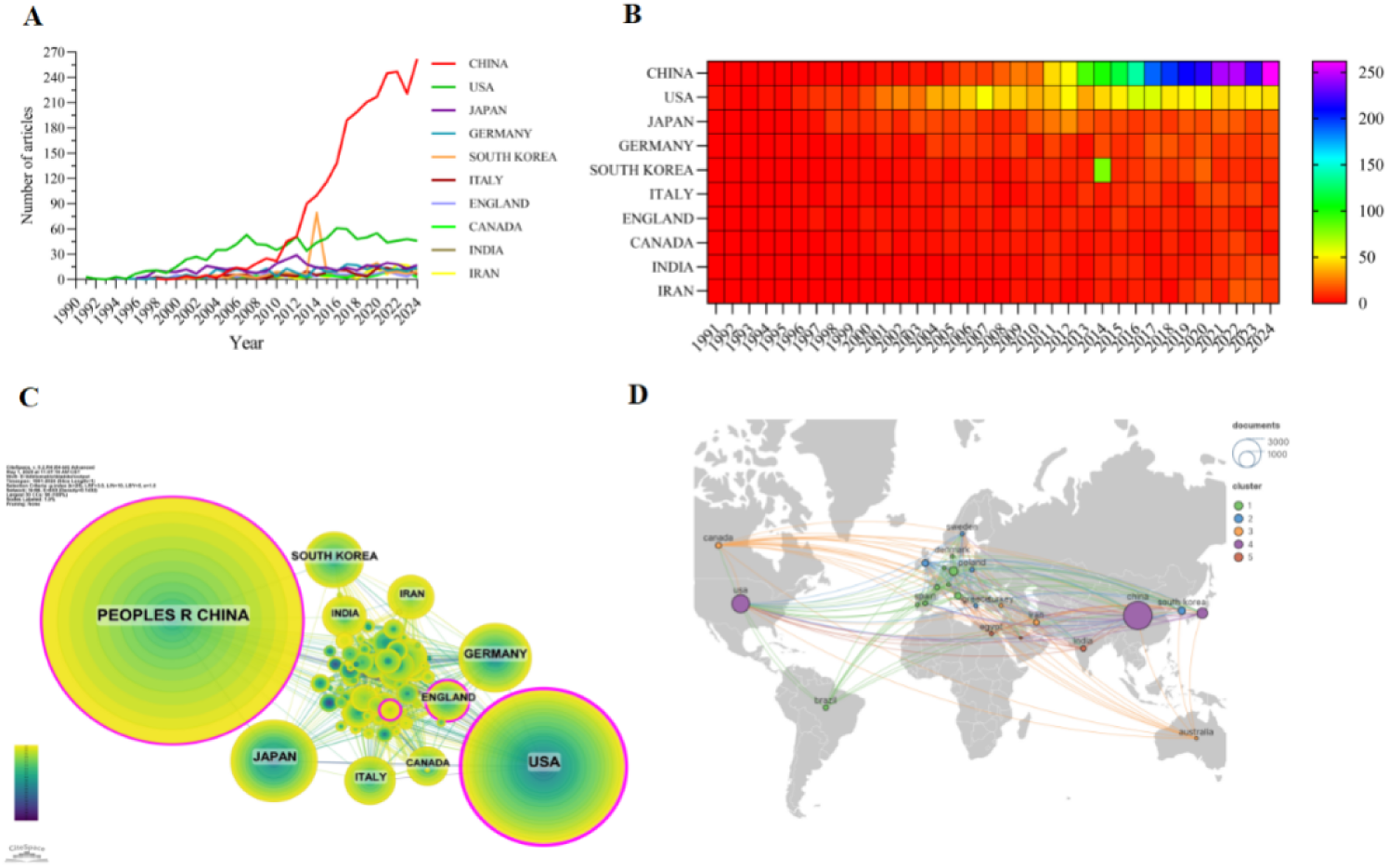
(A). Line graph of national publications (B). Heat map of national publications (C). Networks of country cooperation (D). Knowledge visualization map for international collaboration

**Table 1.** Table of country published literature

| Rank | Country/region | Article counts | centrality | Percentage (%) | Citation | Citation per publication |
| --- | --- | --- | --- | --- | --- | --- |
| 1 | CHINA | 2447 | 0.11 | 45.38% | 62119 | 25.39 |
| 2 | USA | 1093 | 0.5 | 20.27% | 54052 | 49.45 |
| 3 | JAPAN | 390 | 0.05 | 7.23% | 14355 | 36.81 |
| 4 | GERMANY | 256 | 0.09 | 4.75% | 8342 | 32.59 |
| 5 | SOUTH<br>KOREA | 186 | 0.05 | 3.45% | 5657 | 30.41 |
| 6 | ITALY | 165 | 0.06 | 3.06% | 5420 | 32.85 |
| 7 | ENGLAND | 152 | 0.1 | 2.82% | 8151 | 53.63 |
| 8 | CANADA | 120 | 0.08 | 2.23% | 5100 | 42.50 |
| 9 | INDIA | 112 | 0.07 | 2.08% | 2826 | 25.23 |
| 10 | IRAN | 108 | 0.07 | 2.00% | 2436 | 22.56 |

### 3.3 Institutions

A total of 654 institutions have published research on bladder cancer cell death (Table 2; Fig 4). All the 10 institutions by publication volume originated from China and the United States. The University of Texas System led with 178 publications (10,335 citations; average citation = 58.06), followed by UTMD Anderson Cancer Center (2nd; 123 publications, 6,967 citations; average citation = 56.64) and Nanjing Medical University (3rd; 122 publications, 3,689 citations; average citation = 30.02). Further analysis revealed a pronounced preference for intra-country collaboration among both domestic and international institutions. This observed bias highlights the need to strengthen global research partnerships and mitigate academic fragmentation.

**Fig 4.**
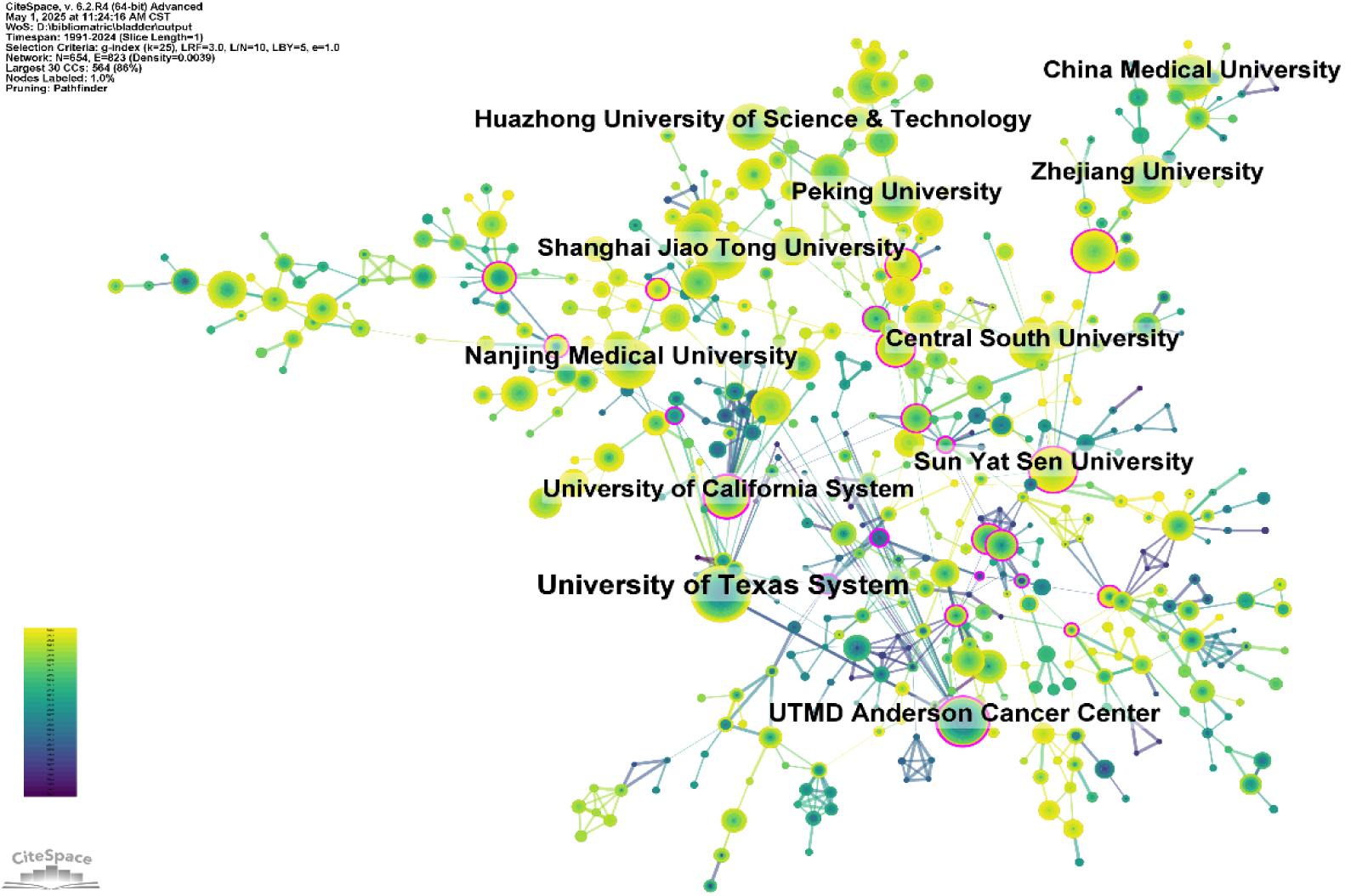
Networks of institutional co-operation

**Table 2.** Table of Institutional Published Literature

| Rank | Institution | Country | Number<br>of studies | Total<br>citations | Average<br>citation |
| --- | --- | --- | --- | --- | --- |
| 1 | University of Texas System | USA | 178 | 10335 | 58.06 |
| 2 | UTMD Anderson Cancer Center | USA | 123 | 6967 | 56.64 |
| 3 | Nanjing Medical University | China | 122 | 3689 | 30.24 |
| 4 | Sun Yat Sen University | China | 105 | 3426 | 32.63 |
| 5 | Zhejiang University | China | 99 | 3440 | 34.75 |
| 6 | Shanghai Jiao Tong University | China | 97 | 2311 | 23.82 |
| 7 | Huazhong University of Science<br>& Technology | China | 97 | 2996 | 30.89 |
| 8 | Peking University | China | 96 | 2525 | 26.30 |
| 9 | China Medical University | China | 93 | 2205 | 23.71 |
| 10 | University of California System | USA | 91 | 5130 | 56.37 |

**Table 3.**
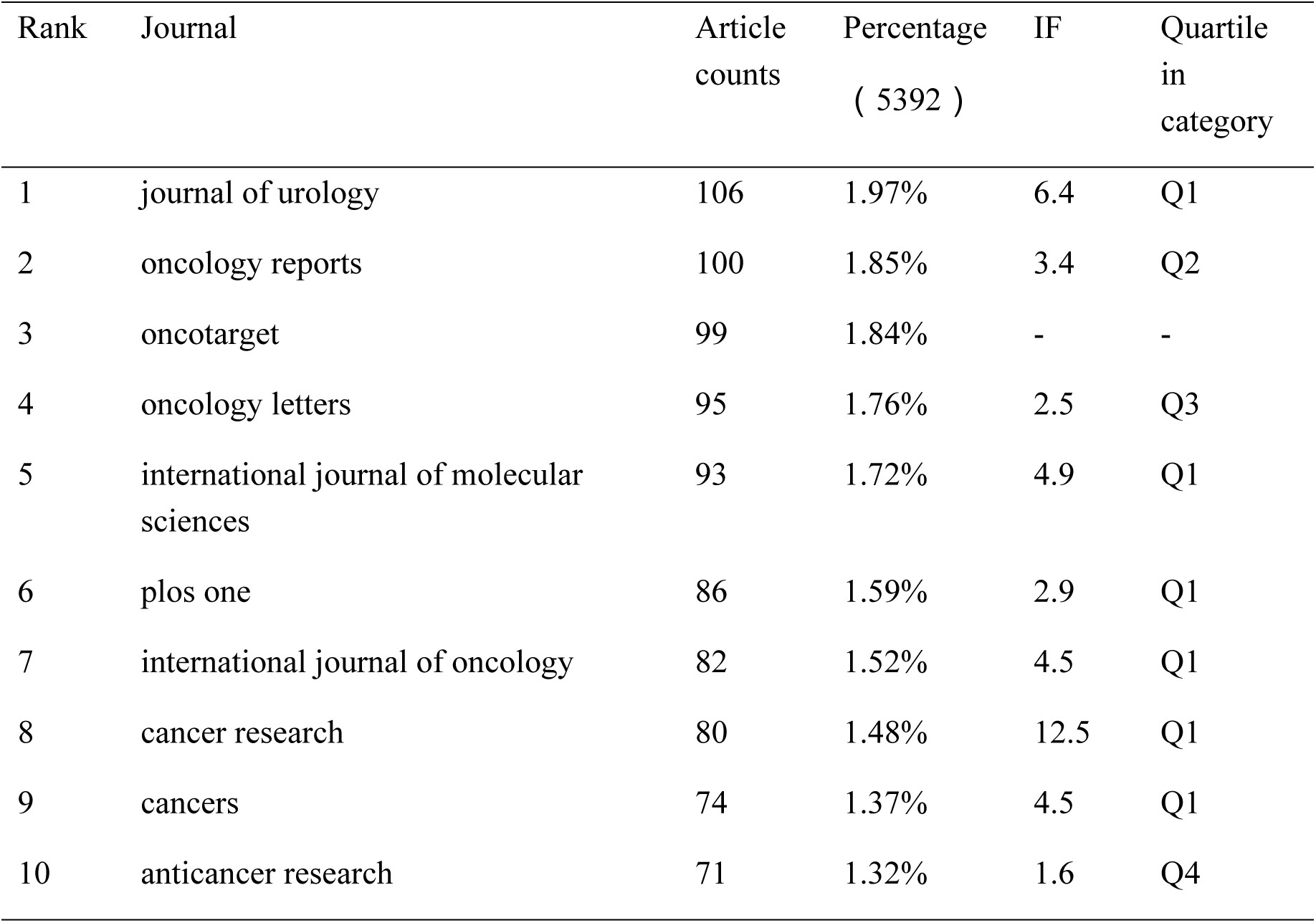
Table of Journal Publications

### 3.4 Journals

<u>Tables 3</u> and <u>Figure4</u> list the top 10 journals by publication volume and co-citation frequency, respectively. The Journal of Urology led in productivity (106 publications, 1.97%), followed by Oncology Reports (100, 1.85%), Oncotarget (99, 1.84%), Oncology Letters (95, 1.76%), International Journal of Molecular Sciences (93, 1.72%), and PLOS ONE (86, 1.59%). Among these, Cancer Research had the highest impact factor (IF 12.5). Notably, 60% were JCR Q1 journals—a dominance attributable to the field’s alignment with high-impact, rigorously peer-reviewed venues, reflecting the topic’s translational significance and interdisciplinary nature. Journal influence was further assessed through co-citation analysis (Fig 5B; Table 4). CANCER RESEARCH ranked first (3,405 co-citations), ahead of ONCOGENE (2,356) and CLINICAL CANCER RESEARCH (2,290). Nature, while 4th in co-citations (2,273), achieved the highest citation density (50.5 citations/article) among top journals. Eighty percent of co-cited journals belonged to JCR Q1, reinforcing their foundational role in the field’s knowledge structure.

**Fig 5.**
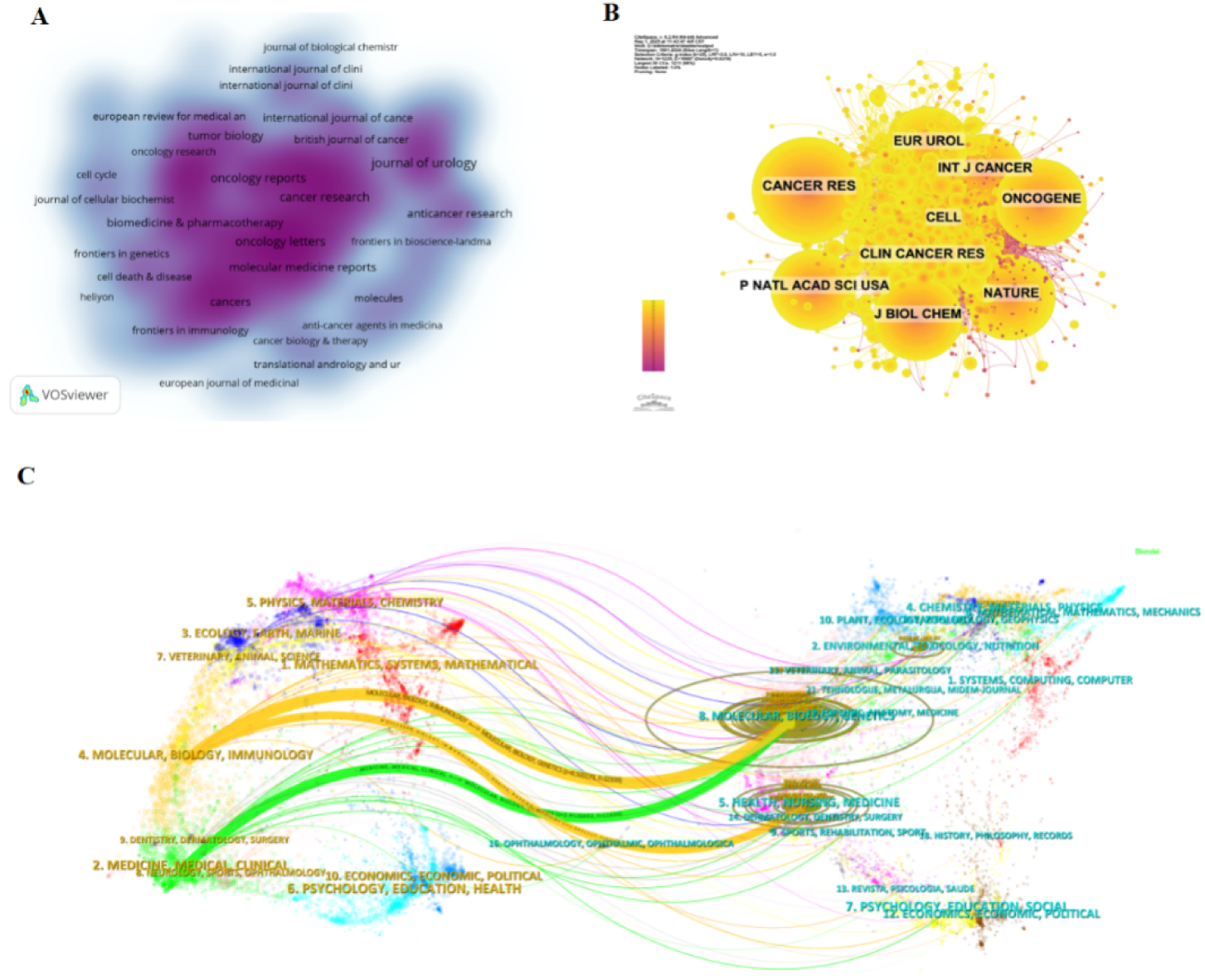
(A). Density map of journal publications (B). Co-citation network map of journals (C). dual map of journals

**Table 4.** Co-citation table of journals

| Rank | Cited Journal | Co-Citation | IF<br>( 2024 ) | Quartile in<br>category |
| --- | --- | --- | --- | --- |
| 1 | CANCER RES | 3405 | 12.5 | Q1 |
| 2 | ONCOGENE | 2356 | 6.9 | Q1 |
| 3 | CLIN CANCER RES | 2290 | 10.4 | Q1 |
| 4 | NATURE | 2273 | 50.5 | Q1 |
| 5 | J BIOL CHEM | 2084 | 4.0 | Q2 |
| 6 | CELL | 2035 | 45.6 | Q1 |
| 7 | P NATL ACAD SCI USA | 2002 | 9.4 | Q1 |
| 8 | INT J CANCER | 1916 | 5.7 | Q1 |
| 9 | EUR UROL | 1844 | 25.3 | Q1 |
| 10 | PLOS ONE | 1798 | 2.9 | Q1 |

The density visualization (Fig 5A) revealed three research clusters: Core oncology: Cancer Research, British Journal of Cancer, International Journal of Cancer Clinical translation: Journal of Urology, Translational Andrology and Urology, Biomedicine & Pharmacotherapy. Basic mechanisms: Journal of Biological Chemistry, Journal of Cellular Biochemistry, Frontiers in Genetics. Dual-map overlay (Fig 5C) traced citation pathways, identifying three key knowledge flows: Yellow trajectory: Publications in Molecular/Biology/Immunology journals influence both basic science (Molecular/Biology/Immunology) and clinical (Health/Nursing/Medicine) domains. Green trajectory: Medicine/Medical Clinical research draws significantly from Health/Nursing/Medicine insights. This topology indicates an evolving research front (left clusters) and established knowledge base (right clusters), predicting future diversification from molecular mechanisms toward clinical and translational oncology.

Dual-map overlay analysis (Fig 5C) identified three citation trajectories: Yellow: Molecular/biology/immunology research influencing both its own domain and health/nursing/medicine; Green: Medicine/clinical publications informed by health/nursing/medicine insights; Predicted shift: Emerging focus from molecular/biology/immunology toward clinical applications. Collectively, these trajectories indicate evolving research fronts in bladder cancer cell death mechanisms."

### 3.5 Authors

Table 5 lists the top 10 most prolific authors in bladder cancer cell death research. Collectively, these authors contributed 273 publications (5.06% of total publications). Kim Wun-Jae led with 37 papers, followed by Liu Yu-Chen and Zhang Wei (33 papers), Choi Yung-Hyun (29 papers), and Chen Wei (25 papers).CiteSpace co-authorship analysis (Fig 6A) revealed distinct research clusters led by prominent authors (e.g., Kim wun-jae, Zhang wei, Liu yuchen), while peripheral contributors exhibited limited collaborative engagement, suggesting more independent research trajectories. Fig 6B and Table 5 further identify the top 10 co-cited and cited authors. The co-citation network’s largest nodes represent the most influential scholars: Jemal a (446 co-citations), Siegel rl. (441), Wang Y. (316), Witjes Ja (315), and Babjuk m (305).

**Fig 6.**
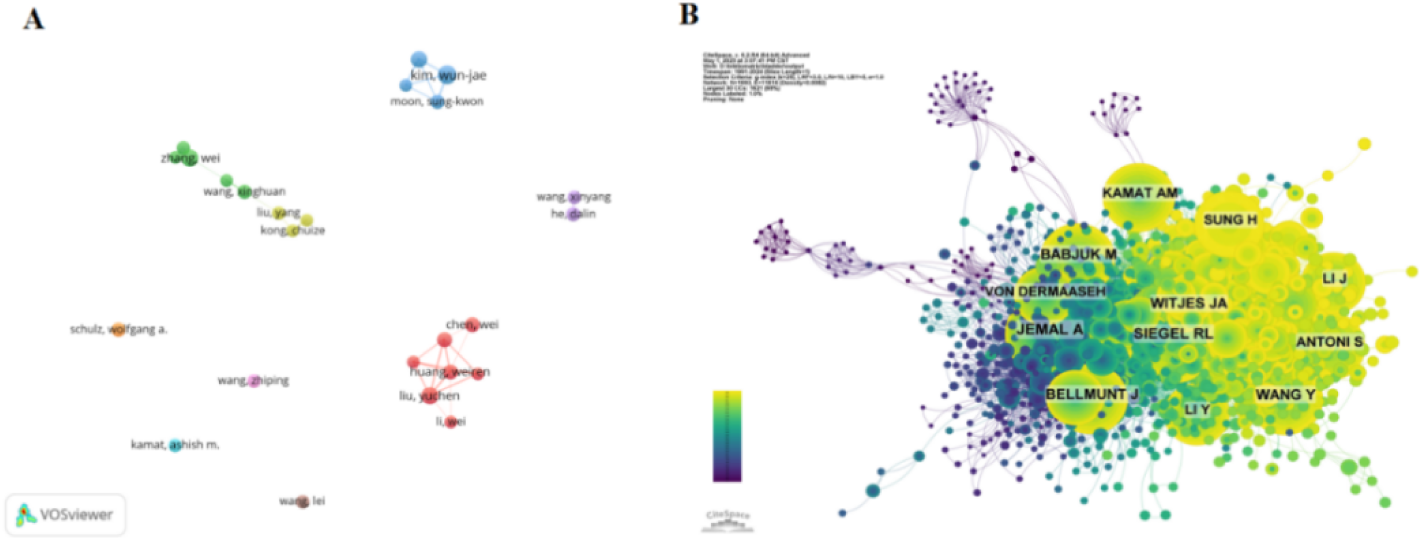
Cooperation network of authors (B). Co-citation network of authors

**Table 5.**
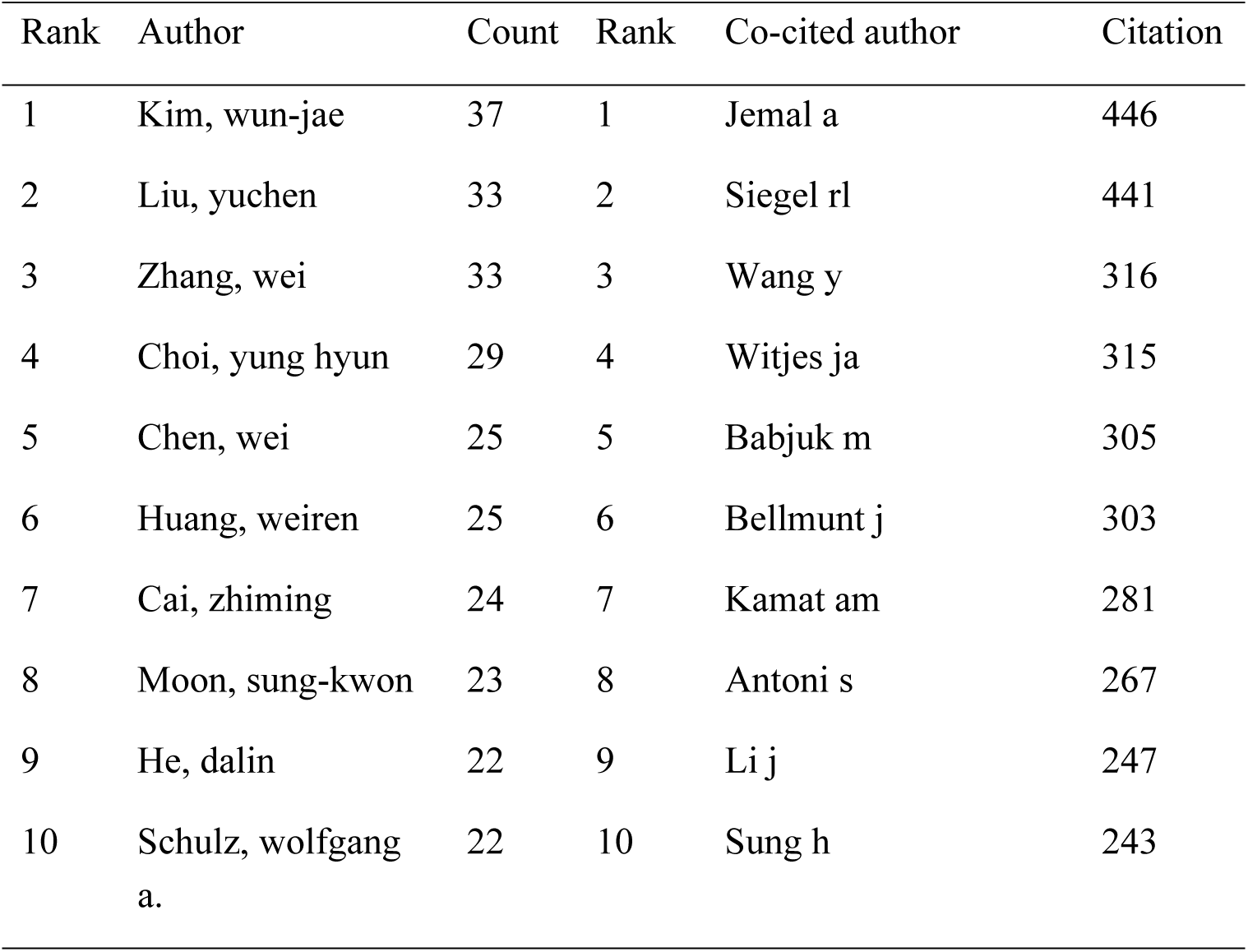
Author’s publications and co-citation table

### 3.6 Citation and co-citation analysis

Citation analysis serves as a critical method for assessing scholarly impact, reflecting both a publication’s influence within a field and prevailing research trends. Table 6 lists the top 10 most-cited papers in bladder cancer cell death research. The highest-cited publication, ’Comprehensive molecular characterization of urothelial bladder carcinoma’ (110 citations), established foundational molecular frameworks for bladder cancer cell death. This was followed by ’Pembrolizumab as Second-Line Therapy for Advanced Urothelial Carcinoma’ (103 citations), which advanced systemic therapy approaches, and ’Atezolizumab in patients with locally advanced and metastatic urothelial carcinoma…’ (79 citations), which validated immune checkpoint inhibition efficacy. Collectively, these seminal works provide conceptual foundations for modern bladder cancer cell death research **Error! Reference source not found.Error! Reference source not found.**.

**Table 6.** Co-citation table of literature

| Rank | Title | Journal | author(s) | Total citations |
| --- | --- | --- | --- | --- |
| 1 | Comprehensive molecular characterization of urothelial bladder carcinoma | NATURE | Weinstein JN | 110 |
| 2 | Pembrolizumab as Second-Line Therapy for Advanced Urothelial Carcinoma | NEW ENGL J MED | Bellmunt J | 103 |
| 3 | Atezolizumab in patients with locally advanced and metastatic urothelial carcinoma who have progressed following treatment with platinum-based chemotherapy: a single-arm, multicentre, phase 2 trial | LANCET | Rosenberg JE | 79 |
| 4 | Comprehensive Molecular Characterization of Muscle-Invasive Bladder Cancer | CELL | Robertson AG | 79 |
| 5 | Advances in bladder cancer biology and therapy | NAT REV CANCER | Tran LD | 78 |
| 6 | Nivolumab in metastatic urothelial carcinoma after platinum therapy (CheckMate 275): a multicentre, single-arm, phase 2 trial | LANCET ONCOL | Sharma P | 66 |
| 7 | Atezolizumab as first-line treatment in cisplatin-ineligible patients with locally advanced and metastatic urothelial carcinoma: a single-arm, multicentre, phase 2 trial | LANCET | Balar AV | 66 |
| 8 | Molecular biology of bladder cancer: new insights into pathogenesis and clinical diversity | NAT REV CANCER | Knowles MA | 60 |
| 9 | MPDL3280A (anti-PD-L1) treatment leads to clinical activity in metastatic bladder cancer | NATURE | Powles T | 50 |
| 10 | Atezolizumab versus chemotherapy in patients with platinum-treated locally advanced or metastatic urothelial carcinoma (IMvigor211): a multicentre, open-label, phase 3 randomised controlled trial | LANCET | Powles T | 49 |

To elucidate intellectual relationships within the citation landscape, we performed co-citation clustering analysis on 5,392 publications (1991-2024) using time-slicing. This generated a comprehensive co-citation network (Fig 7A) comprising 2,046 nodes and 8,431 links (density = 0.004). The analysis identified 19 distinct clusters (Figs 7B and 7C), ranked by size from largest (0) to smallest (19). The network demonstrated exceptional structural validity, with high modularity (Q = 0.8388) and mean silhouette score (S = 0.8817), confirming robust cluster homogeneity.These co-citation clusters represent the intellectual foundation of bladder cancer cell death research. We synthesized five major research hotspots from the 19 clusters: (1) Cell Death Mechanisms & Molecular Regulation: 0 Ferroptosis, 1 Bcl-2, 2 Survivin; (2) Bladder Cancer Phenotypes: 3 Urothelial carcinoma, 16 Bladder cancer transformation; (3) Therapeutic Interventions: 4 Proliferation pathways, 7 Ursolic acid, 11 Metformin, 9 Allyl isothiocyanate. (4) Biomarkers & Diagnostics: 8 Tumor markers, 5 lncRNAs, 6 miRNAs, 12 Circular RNAs. (5) Clinical Translation: 10 Animal models, 13 Clusterin, 14 FAS-ligand, 17 Clinical implications, 18 ROS signaling.

**Fig 7.**
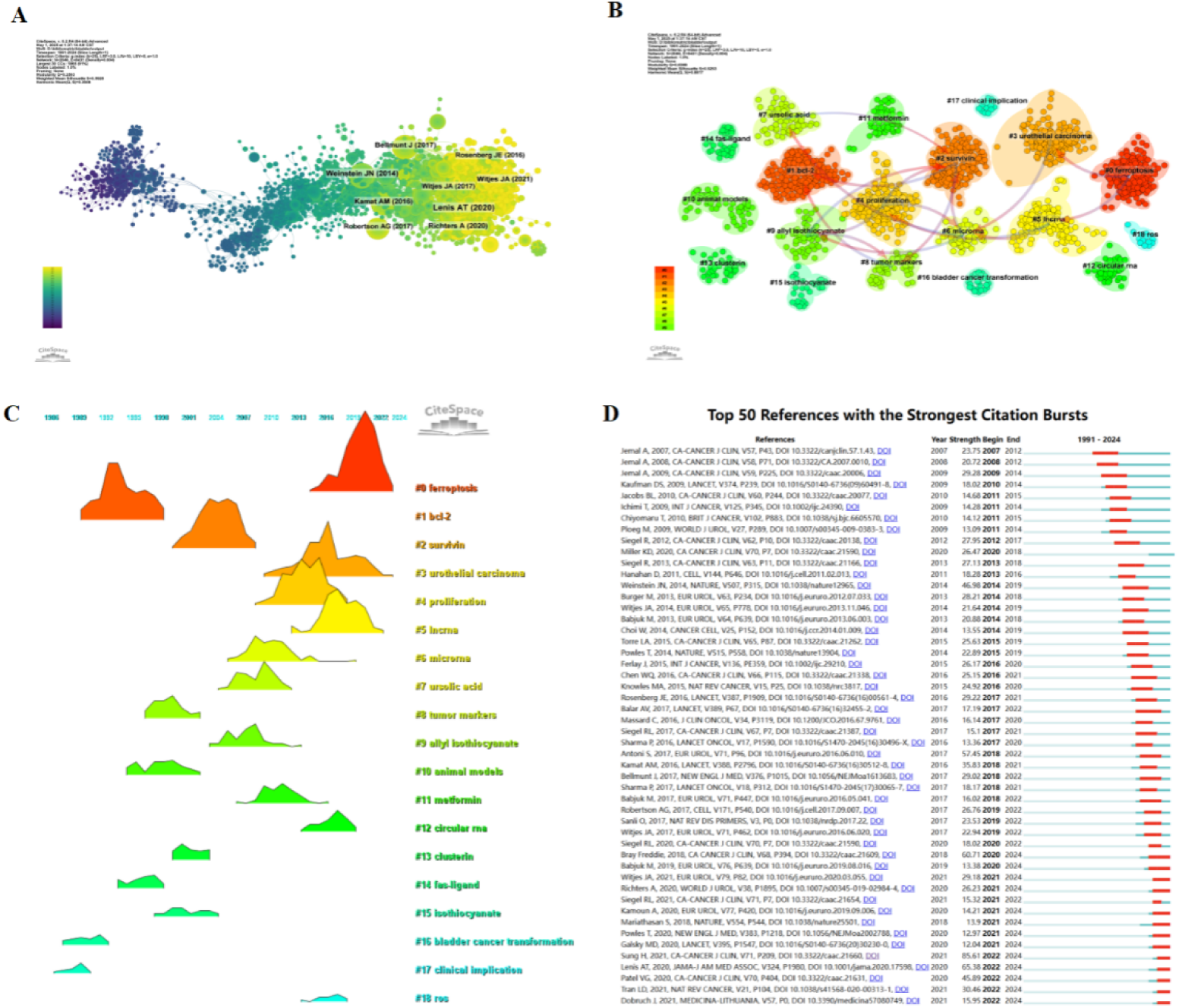
co-cited network of literature (B). Clustering of co-cited literature (C). Peak map of co- cited literature (D). Bursting map of cited literature

Citation burst analysis reveals rapidly evolving research frontiers. Fig 7D identifies the top 50 references with the strongest citation bursts. The earliest significant burst (2007) emerged from seminal work by Jemal et al., which highlighted bladder cancer mortality rates and galvanized research interest in therapeutic development. The most intense burst (strength = 85.61) was observed for Sung et al.’s 2021 CA: A Cancer Journal for Clinicians study, persisting through 2024.Current citation bursts (initiated 2022 and ongoing) are driven by five influential publications: Sung H, et al. (2021) CA Cancer J Clin **Error! Reference source not found.**, Lenis AT, et al. (2020) JAMA**Error! Reference source not found.**, Patel VG, et al. (2020) CA Cancer J Clin **Error! Reference source not found.**, Tran L, et al. (2020) Nature **Error! Reference source not found.**, Dobruch J, et al. (2021) Medicina (Kaunas) **Error! Reference source not found.**. Collectively, these bursting publications—dominated by epidemiological updates and clinical guideline revisions—demonstrate the field’s dynamic evolution toward translational applications.

### 3.7 Keywords

Keywords encapsulate core research concepts, and their co-occurrence analysis reveals both disciplinary hotspots and emerging trends. Using VOSviewer, we constructed a keyword co-occurrence network from 5,392 publications (Table 7; Figs 8A and 8B). The most frequent terms were apoptosis (2,299 occurrences), bladder cancer (2,045), proliferation (797), growth (570), and activation (500).Network clustering identified five thematic domains: Cell Death Mechanisms (Green): apoptosis, autophagy, mitochondrial dysfunction, ROS; Tumor Progression (Blue): bladder cancer, proliferation, migration, metastasis; Clinical Therapeutics (Red): chemotherapy, targeted therapy, doxorubicin, cisplatin, PD-1/PD-L1 inhibition; Molecular Biomarkers (Yellow): p53, biomarkers; Disease Subtyping (Purple): urothelial carcinoma.

**Fig 8.**
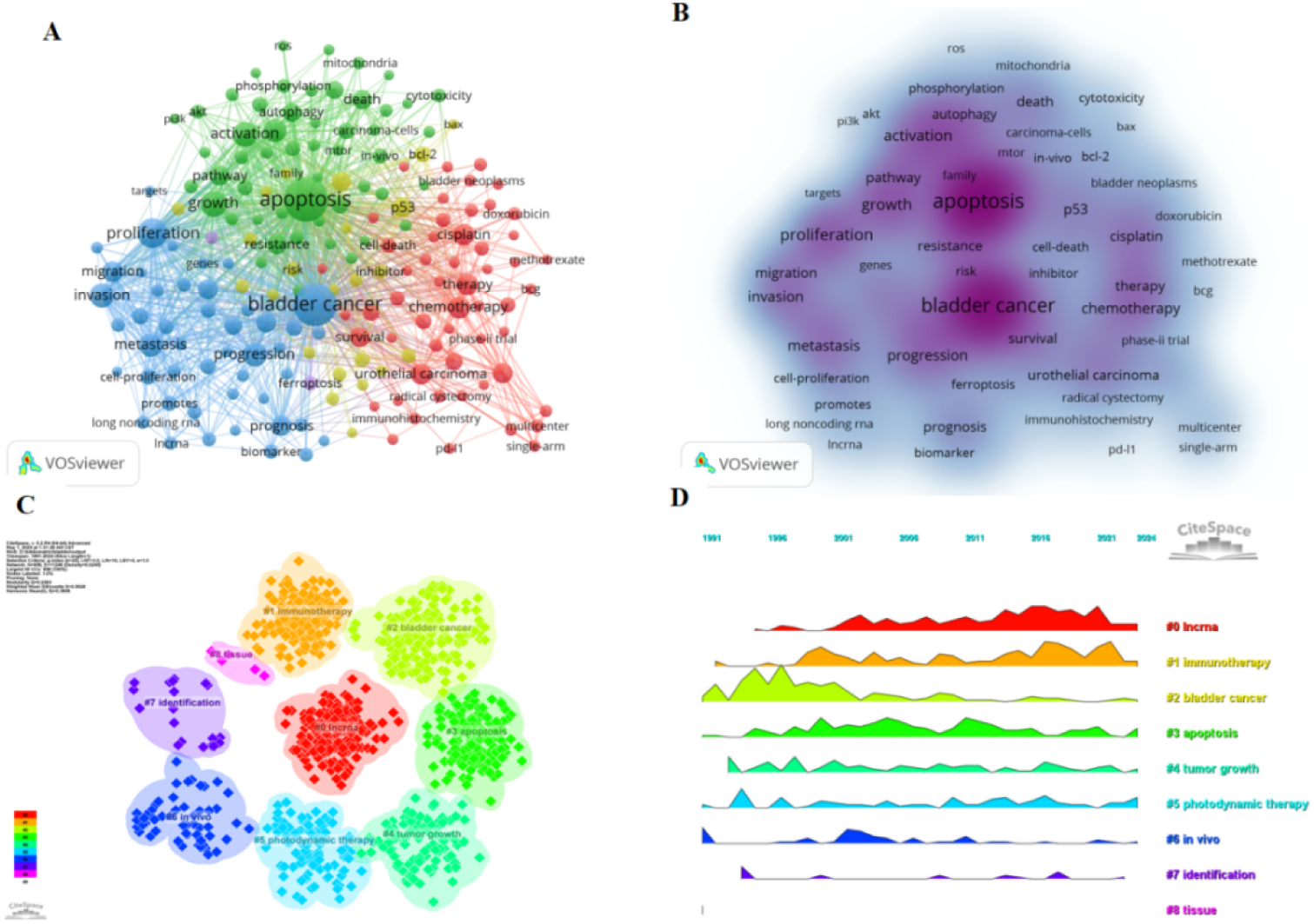
(A). Network map of high-frequency keywords (B). Density map of keywords (C). Peak map of keyword clustering (D). clustering map of keywords

**Table 7.**
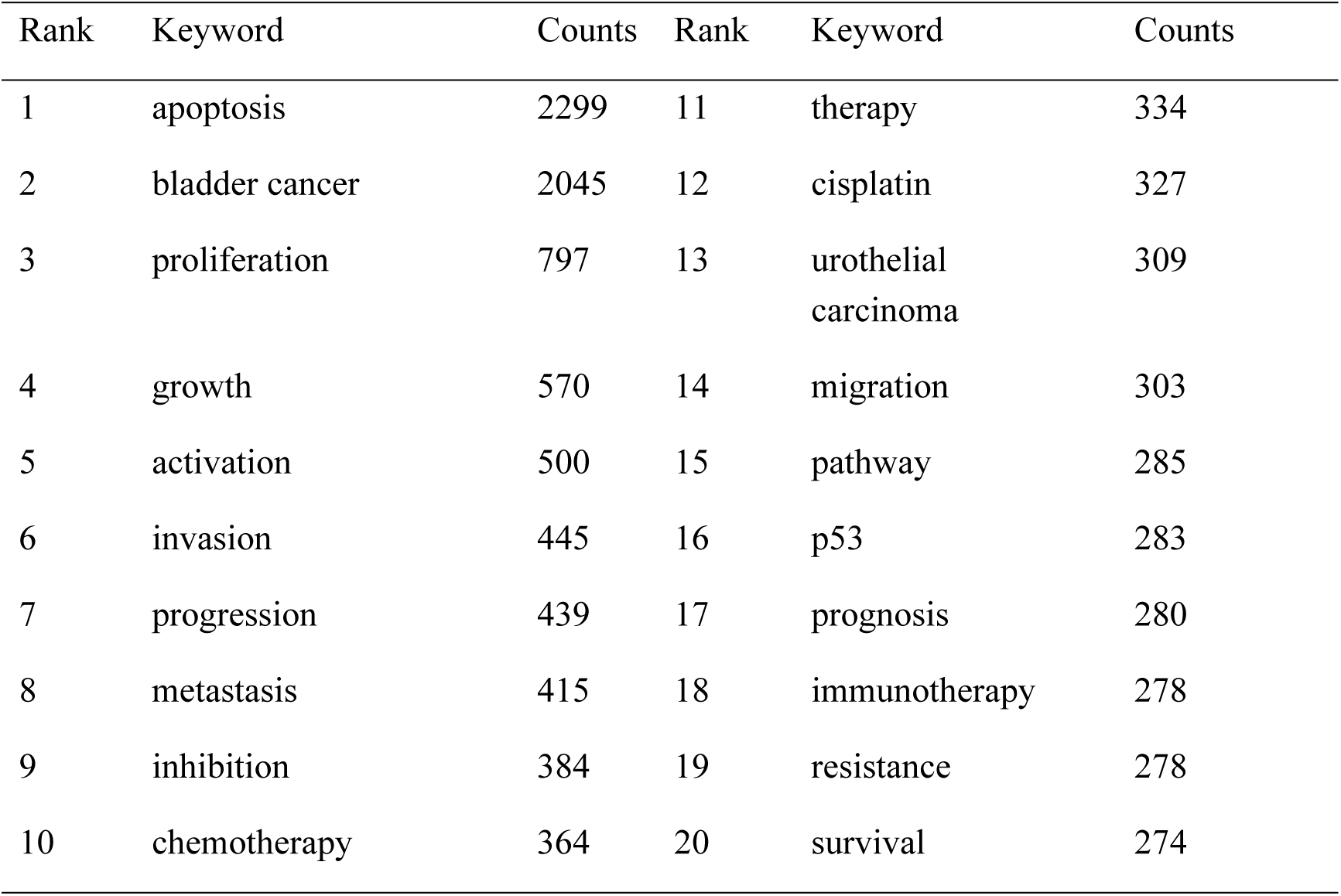
High Frequency Keyword Table

| Rank | Keyword | Counts | Rank | Keyword | Counts |
| --- | --- | --- | --- | --- | --- |
| 1 | apoptosis | 2299 | 11 | therapy | 334 |
| 2 | bladder cancer | 2045 | 12 | cisplatin | 327 |
| 3 | proliferation | 797 | 13 | urothelial carcinoma | 309 |
| 4 | growth | 570 | 14 | migration | 303 |
| 5 | activation | 500 | 15 | pathway | 285 |
| 6 | invasion | 445 | 16 | p53 | 283 |
| 7 | progression | 439 | 17 | prognosis | 280 |
| 8 | metastasis | 415 | 18 | immunotherapy | 278 |
| 9 | inhibition | 384 | 19 | resistance | 278 |
| 10 | chemotherapy | 364 | 20 | survival | 274 |

Using CiteSpace, we generated timeline visualizations (Figs 8C and 8D) to illustrate the temporal evolution of research hotspots. Our analysis identified lncRNA, bladder cancer, apoptosis, tumor growth, photodynamic therapy, in vivo studies, identification, and tissue analysis as current predominant research foci. Furthermore, we performed keyword burst analysis via CiteSpace, selecting the top 50 terms with the strongest citation bursts during 1991–2024, ranked by burst intensity (Fig 9). These terms represent the field’s current research priorities and potential future directions. Notably, "invasion" exhibited the highest burst strength. Given that invasion underpins the entire progression, metastasis, and prognosis of bladder cancer, it constitutes a key entry point for understanding malignant behavior and optimizing diagnosis/therapy. Additionally, keywords including "pembrolizumab", "multicenter" trials, "metabolism", the "tumor microenvironment", "cell death" mechanisms, and "long noncoding RNA" (lncRNA) continued to demonstrate significant burst strength in 2024, strongly suggesting these areas will likely remain central to future research.

**Fig 9.**
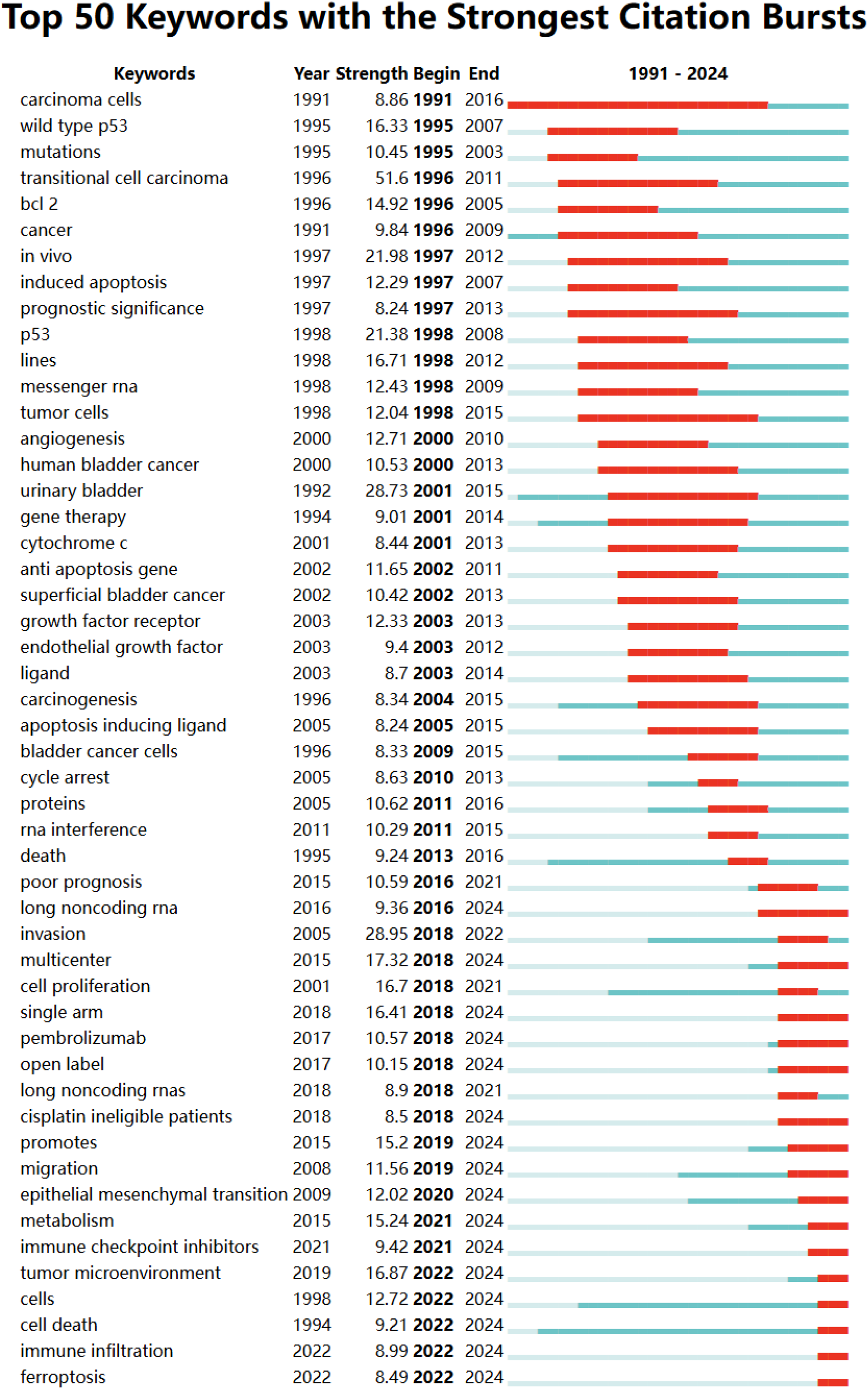
Bursting map of keywords

## 4 Discussion

### 4.1 General information

A comprehensive search of the WoSCC database (1991–2024) identified 5,392 publications on bladder cancer cell death mechanisms, contributed by 1,693 authors from 654 institutions across 95 countries/regions and published in 1,235 journals. Despite intermittent fluctuations, annual publication output exhibited an overall upward trajectory (Fig 2). Notably, a sharp surge occurred after 2016, peaking significantly in 2017. Citation burst analysis (Fig 7D) attributed this peak to increased research focus on programmed cell death-based chemotherapy for bladder cancer patients beginning in 2017. Specifically, 7 of the top 50 references with the strongest citation bursts were published in 2017, including 3 directly addressing chemotherapy. Leading contributors Balar AV, Sharma P, and Bellmunt J investigated chemotherapy regimens targeting programmed cell death (using agents including atezolizumab, pembrolizumab, and nivolumab), thereby establishing the foundation for subsequent research on cell death mechanism-based immunotherapy combinations in bladder cancer **Error! Reference source not found.Error! Reference source not found.**. Concurrently, geographic collaboration networks revealed the top 10 contributing countries spanning three regions: Asia, Europe, and North America. Of these, China emerged as a dominant contributor, ranking among the top three in both publication volume and citation frequency. The sustained increase in annual publications and citation frequency since 1991 reflects intensifying global interest in bladder cancer cell death research, underscoring its growing academic significance and translational potential.

An in-depth analysis of contributions from 95 countries/regions revealed a heterogeneous global research landscape. China emerged as the dominant contributor in bladder cancer cell death research, evidenced by its leading publication output (2,447 papers, 45.38% of total) and high citation frequency (62,199 citations). Notably, China is not only the sole developing nation among the top 10 contributing countries but has also demonstrated exponential publication growth since 2011, establishing a significant lead over other nations. While China dominates in research volume, the United States and United Kingdom lead in citation impact (mean citations: 49.45 and 53.63, respectively), reflecting higher-quality contributions. This disparity underscores the critical need for enhanced international collaboration to simultaneously augment the volume and quality of global research output **Error! Reference source not found.**.

Institutional contributions reflect distinct national research trajectories in this field. Notably, 7 of the top 10 institutions by publication volume are in China, while the remaining 3 are in the United States. Among these, the University of Texas System, UTMD Anderson Cancer Center, and Nanjing Medical University lead bladder cancer cell death mechanism research, contributing substantially to global output. Critically, Chinese institutions comprise 70% of the top 10 contributors and account for 45.38% of total publications, underscoring their dominant quantitative role. Conversely, U.S. institutions exhibit significantly higher per-publication citation rates (49.45 vs. China’s 25.39), reflecting greater emphasis on high-impact research. Furthermore, the three most-cited institutions—University of Texas System (10,335 citations), UT MD Anderson Cancer Center (6,967), and University of California System (5,130)—far surpass leading Chinese institutions (e.g., Nanjing Medical University: 3,689; Sun Yat-sen University: 3,426). This citation disparity may stem from language barriers, data-sharing policies, and divergent research priorities. Specifically, U.S. institutions (e.g., University of Texas System) maintain broader international collaborations, publishing predominantly in high-impact English journals; in contrast, Chinese institutions focus on domestic networks, indicating opportunities for enhanced global engagement. To address these gaps, prioritized strategies include: (1) establishing multinational consortia for data sharing; (2) developing open-data platforms compliant with privacy regulations; and (3) overcoming language barriers via professional translation services and standardized English scientific reporting. Implementing these measures will amplify global impact and accelerate clinical translation of bladder cancer cell death research.

Influential authors including Jemal A, Bellmunt J, Witjes JA, and Siegel RL have profoundly shaped this field through their work on bladder cancer epidemiology, cell death mechanisms, and therapeutic applications. Notably, Bellmunt J pioneered research on programmed cell death-based therapies for BC **Error! Reference source not found.**, establishing him as a key opinion leader. Concurrently, Witjes JA’s clinical studies on NMIBC treatment **Error! Reference source not found.** directly informed subsequent clinical decisions regarding cell death mechanism-immunotherapy synergies. These contributions underscore the critical role of foundational research in driving therapeutic innovation. Regarding dissemination, Journal of Urology, Oncology Reports, and Oncotarget led in publication volume for bladder cancer cell death research. However, citation impact was dominated by high-impact journals: CANCER RESEARCH (IF=12.5; co-citations: 3,405), NATURE (IF=50.5; co-citations: 2,273), and CELL (IF=45.6; co-citations: 2,035). Strikingly, 90% of the top 10 co-cited journals (e.g., European Urology, ONCOGENE) are JCR Q1 publications, emphasizing the field’s academic significance. Coupling analysis further revealed that key findings are primarily published in immunology, molecular biology, and genetics journals. Cross-disciplinary integration of fundamental sciences— particularly molecular biology, genetics, biomaterials, and medicinal chemistry— provides the essential framework for elucidating cell death mechanisms and translating discoveries into clinical applications. Consequently, multidisciplinary collaboration across these domains is poised to accelerate therapeutic optimization and advance precision medicine for bladder cancer patients.

### 4.2 Frontiers

Co-citation networks mapped via CiteSpace and VOSviewer reveal distinct thematic clusters representing major research foci in bladder cancer cell death (Fig 8). These span a multidimensional continuum from fundamental mechanisms to clinical translation, with hotspots concentrated in: Cell death mechanisms (apoptosis, autophagy, mitochondrial dysfunction, ROS signaling); Tumor progression (bladder carcinogenesis, proliferation, migration, metastasis); Clinical strategies (chemotherapy, targeted therapy, immunohistochemical biomarkers). Critically, immunotherapy has emerged as a dominant clinical paradigm. Contemporary trials position immune-based therapies as transformative interventions—specifically through their capacity to reprogram the tumor microenvironment (TME)—and dominate treatment research by leveraging cell death pathways for antitumor efficacy **Error! Reference source not found.Error! Reference source not found.**. This predominance underscores the therapeutic imperative of cell death mechanism-based immunotherapy in bladder cancer.

PCD mechanisms—including apoptosis and ferroptosis—represent a critical research frontier in bladder oncology, profoundly influencing tumorigenesis, progression, prognostic prediction, and therapeutic interventions**Error! Reference source not found.Error! Reference source not found.**. Therefore, current research focuses on multi-faceted breakthroughs targeting these core mechanisms. Specifically, in targeted drug delivery, nanoparticle-based systems demonstrate substantial potential for overcoming traditional therapeutic limitations **Error! Reference source not found.Error! Reference source not found.**. These systems enhance solubility, tumor permeability, and targeting specificity of PCD-inducing agents, thereby reducing required dosages and systemic toxicity while precisely augmenting tumor cell death induction, which improves therapeutic precision and efficacy. Concurrently, the TME receives considerable attention due to its pivotal regulatory role in PCD. Complex TME interactions—such as immunosuppression and therapy resistance induction—coupled with its secreted growth factors, cytokines, and chemokines, not only support tumor growth and metastasis but also significantly impair therapeutic PCD induction. Consequently, elucidating and modulating TME-mediated PCD regulation emerges as the key strategy to overcome treatment resistance**Error! Reference source not found.**.

Co-citation analysis of research trends highlights the evolving focus bladder cancer cell death mechanisms. Initially, early studies emphasized fundamental roles of cell death, demonstrating that tumor suppressor p53 critically regulates cell cycle progression and apoptosis under cytotoxic conditions **Error! Reference source not found.Error! Reference source not found.**. Subsequently, mid-phase research explored therapeutic applications, revealing synergistic effects between chemotherapy/radiotherapy and cell death pathways. Chemotherapeutic agents were confirmed to eliminate bladder cancer cells via apoptosis; unfortunately, tumor cells frequently developed chemoresistance. Consequently, researchers adopted combination strategies—such as chemo-radiotherapy—to induce cell death through complementary mechanisms, significantly enhancing treatment efficacy **Error! Reference source not found.Error! Reference source not found.**. Recently, investigations shifted toward precision medicine, integrating immunotherapy and PDT with cell death mechanisms **Error! Reference source not found.Error! Reference source not found.**. Collectively, this progression reflects the fields dynamism and its continuous advancement toward technologically driven clinical solutions.

#### 4.2.1 Cell Death Mechanisms in Bladder Cancer: an in-depth exploration of bladder cancer

PCD constitutes an active cellular demise process triggered by specific signals to maintain internal homeostasis. Major PCD modalities include ferroptosis, apoptosis, cuproptosis, necroptosis, autophagy, and pyroptosis **Error! Reference source not found.Error! Reference source not found.**. Notably, apoptosis and ferroptosis represent critical anti-cancer mechanisms. Bladder cancer cells frequently evade elimination by acquiring apoptosis resistance, enabling survival, proliferation, and metastasis. Thus, apoptosis evasion constitutes a major therapeutic challenge. Critically, studies confirm that Survivin, a pivotal anti-apoptotic protein, directly mediates chemoresistance in bladder cancer. Therapeutic suppression of Survivin expression significantly enhances treatment efficacy while potently reducing chemoresistance development in preclinical models **Error! Reference source not found.Error! Reference source not found.**. Furthermore, the B-cell lymphoma 2 (BCL-2) gene encodes a mitochondrial protein that regulates apoptotic PCD **Error! Reference source not found.**, while tumor suppressor p53 aberrantly influences bladder cancer chemoresistance through apoptosis induction **Error! Reference source not found.**. Collectively, these dysregulated factors (Survivin, BCL-2, p53) not only drive tumor progression but also serve as poor prognostic indicators **Error! Reference source not found.**. Consequently, targeting apoptotic pathways emerges as a promising clinical strategy for overcoming drug resistance, particularly in advanced or refractory bladder cancer.

Ferroptosis—an iron-dependent, lipid peroxidation-driven form of regulated cell death—is closely associated with bladder cancer progression **Error! Reference source not found.**. Mechanistically, intracellular labile iron (Fe²⁺) catalyzes ROS generation via the Fenton reaction, oxidizing polyunsaturated fatty acids (PUFAs) in cell membranes. This initiates lipid peroxidation cascades, increasing membrane fragility and triggering cell demise. Moreover, impaired glutathione peroxidase 4 (GPX4) activity—a key antioxidant enzyme in the glutathione system—reduces antioxidant capacity, leading to lethal ROS accumulation **Error! Reference source not found.Error! Reference source not found.**. Research indicates that expression levels of ferroptosis suppressor genes, such as GPX4 and SLC7A11, are negatively correlated with tumor malignancy in bladder cancer tissue. Erastin is a classic ferroptosis inducer that inhibits the activity of the key intracellular transporter SLC7A11, thereby significantly reducing the cell’s glutathione (GSH) synthesis capacity, triggering lipid peroxidation and ultimately leading to ferroptosis **Error! Reference source not found.Error! Reference source not found.**. Given ferroptosis plays a critical role in bladder cancer initiation, progression, and therapeutic resistance, its induction has emerged as a promising strategy to trigger cancer cell death, particularly in aggressive malignancies resistant to conventional therapies **Error! Reference source not found.**. Therefore, therapeutic strategies targeting ferroptosis offer novel avenues for precision medicine in bladder cancer.

Substances released during apoptosis and ferroptosis can profoundly alter the composition and function of the TME. Specifically, when ferroptosis occurs in bladder cancer cells, it releases damage-associated molecular patterns (DAMPs), including ATP, high-mobility group box 1 (HMGB1), calreticulin (CRT), and pro-inflammatory cytokines. Notably, HMGB1 can bind receptors on dendritic cells (DCs) within the TME, promoting DC maturation. Mature DCs efficiently engulf, process, and present tumor antigens, thereby facilitating T cell priming and ultimately enhancing tumor cell killing while inhibiting tumor growth **Error! Reference source not found.Error! Reference source not found.**. In contrast, apoptotic cells also release diverse signaling molecules that influence the TME. Apoptotic signaling may affect intratumoral cell competition by stimulating proliferation of neighboring cells and exerting paracrine effects. Furthermore, apoptosis promotes the recruitment of immunosuppressive cells—such as regulatory T cells (Tregs), M2-polarized macrophages, and myeloid-derived suppressor cells (MDSCs)—which can contribute to cancer progression and therapy resistance **Error! Reference source not found.**. Collectively, these findings demonstrate that ferroptosis, apoptosis, and the TME are not isolated entities in bladder cancer pathogenesis. Instead, they interact through intricate molecular mechanisms to co-regulate tumor initiation, progression, metastasis, and therapeutic responses, forming a dynamic, interconnected regulatory network.

#### 4.2.2 Clinical Translation of Cell Death Mechanisms

Translating cell death mechanisms into clinical strategies represents a promising frontier in BC treatment. BC, characterized by high mutational burden, is particularly amenable to immunotherapy, especially checkpoint inhibitors targeting PD-1 or its ligand PD-L1. The interaction between programmed death-ligand 1 (PD-L1) on tumor cells and programmed cell death protein 1 (PD-1) on immune cells critically enables tumor immune evasion. Blocking this interaction with immune checkpoint inhibitors (ICIs) provides a potent strategy for targeted cancer immunotherapy **Error! Reference source not found.Error! Reference source not found.**. Notably, both apoptosis and ferroptosis—when accompanied by intact damage-associated molecular pattern (DAMP) release—can trigger robust immunogenicity. Chemotherapy-induced immunogenic cell death (ICD), which involves DAMP release (e.g., calreticulin, ATP, HMGB1), exemplifies this synergy and enhances immune activation when combined with ICIs **Error! Reference source not found.**. Consequently, combining ferroptosis inducers with ICIs demonstrates synergistic efficacy by enhancing anti-tumor immunity and overcoming immune evasion. Simultaneously, PDT has emerged as an attractive modality in cancer treatment. PDT induces apoptosis or necrosis primarily through ROS-mediated damage to cellular components (e.g., proteins, lipids, nucleic acids) **Error! Reference source not found.**. Crucially, for NMIBC—which predominantly affects the bladder mucosa—intravesical PDT enables precise targeting. Following instillation of a photosensitizer, laser activation at tumor-specific wavelengths selectively eradicates mucosal lesions while minimizing damage to the muscularis propria and urethra, thereby reducing recurrence rates **Error! Reference source not found.**. Importantly, PDT-generated ROS concurrently trigger multiple cell death pathways: high-concentration ROS directly disrupt mitochondrial membranes to initiate apoptosis, while lipid membrane oxidation induces ferroptosis **Error! Reference source not found.Error! Reference source not found.**. Collectively, treatment strategies integrating cell death mechanisms (e.g., with IT or PDT) are gaining significant traction in BC management, creating novel therapeutic possibilities.

### 4.3 Limitations

This study employed bibliometric methods to analyze global research trends and emerging frontiers in bladder cancer cell death mechanisms. Notably, several limitations should be acknowledged. First, our analysis relied exclusively on English-language articles and reviews retrieved from WoSCC. This single-database approach may exclude relevant studies indexed in other databases (e.g., Scopus, PubMed) or published in other languages, potentially limiting the comprehensiveness of our findings. Future studies should integrate data from multiple databases to ensure more comprehensive coverage. Second, publication timelines inherently affect citation metrics; consequently, recent high-impact publications may be underrepresented due to their lower cumulative citation counts. Furthermore, our analysis did not distinguish between bladder cancer subtypes—such as urothelial carcinoma UC and non-urothelial carcinoma—or between MIBC and NMIBC. This broad categorization may obscure subtype-specific research trends. Although these limitations exist, the trends and insights elucidated in this study establish a solid foundation for understanding the current research landscape and guiding future investigations in this rapidly evolving field.

## 5 Conclusion

This bibliometric analysis comprehensively delineates the research landscape of bladder cancer cell death mechanisms from 1991 to 2024, identifying global trends, collaborative networks, and emerging frontiers. Key research foci include apoptosis regulation, ferroptosis pathways, immunotherapy applications, photodynamic therapy, and nanomedicine integration. Critically, our findings underscore the pivotal role of targeting cell death mechanisms to overcome therapy resistance and advance precision oncology in bladder cancer. Future investigations should prioritize exploring the interaction between cell death pathways and the tumor microenvironment, deciphering resistance mechanisms, and formulating targeted therapeutic strategies. These directions are pivotal for driving advancements in bladder cancer treatment and improving patient outcomes. Although constrained by acknowledged limitations, this study provides valuable insights into the evolving research trajectory and translational potential of targeting cell death mechanisms for bladder cancer management.

## Abbreviation

WoSCC: Web of Science Core Collection IT: immunotherapy
PDT: photodynamic therapy BC: Bladder cancer
UC: Urothelial carcinoma
NMIBC: non-muscle-invasive bladder cancer MIBC: muscle-invasive bladder cancer
TURBT: transurethral resection of bladder tumor BCG: Bacillus Calmette-Guérin
PCD: programmed cell death
TME: tumor microenvironment
BCL-2: B-cell lymphoma 2
PUFAs: polyunsaturated fatty acids
GPX4: glutathione peroxidase 4
GSH: glutathione
DAMPs: damage-associated molecular patterns
HMGB1: high-mobility group box 1
DCs: dendritic cells
Tregs: T cells
MDSCs: myeloid-derived suppressor cells
PD-L1: programmed death-ligand 1
PD-1: programmed cell death protein 1
ICIs: immune checkpoint inhibitors
DAMP: damage-associated molecular pattern
ICD: immunogenic cell death
ROS: reactive oxygen species

## Funding

The author(s) declare that financial support was received for the research and/or publication of this article. Key Research and Development Program of Ningxia Hui Autonomous Region (2022BEG03133); Science and Technology Benefiting the People Project of Ningxia Hui Autonomous Region (2024CMG03004); Natural Science Foundation Project of Ningxia (2023AAC02076); Science and Technology Benefiting the People Project of Ningxia Hui Autonomous Region (2022CMG03030).

## Notes

### Competing Interest Statement

The authors have declared no competing interest.

